# CLASP1/2 REGULATE IMMUNE SYNAPSE MATURATION IN NATURAL KILLER CELLS

**DOI:** 10.1101/2025.01.20.633904

**Authors:** Alejandro P. Pariani, Victoria Huhn, Tomás Rivabella Maknis, Victoria Alonso, Evangelina Almada, Rodrigo Vena, Cristián Favre, James R. Goldenring, Irina Kaverina, M. Cecilia Larocca

## Abstract

Natural killer (NK) cells are the first line of defense against viral infections and tumors. Their cytotoxic activity relies on the formation of an immune synapse (IS) with target cells. The lymphocyte function–associated antigen (LFA)-1 plays a central role in NK cell cytotoxicity by modulating NK-IS assembly and maturation. LFA-1 organization at the IS involves a Golgi-dependent mechanism, which has not been completely elucidated. CLIP-associating proteins (CLASP) 1/2 are microtubule plus-tip interacting proteins that control the dynamics of Golgi derived microtubules (GDMTs). In the present study, we found that CLASP1/2 depletion impaired LFA-1 organization at the IS and inhibited the polarization of the centrosome and the lytic granules towards the target cell. Our results also revealed the role of the Golgi apparatus as a microtubule organizing center (MTOC) in these cells. Furthermore, we found that, similarly to what was described in other cell types, NK cells require CLASP1/2 and AKAP350 for efficient nucleation of microtubules at the Golgi. Overall, this study uncovers the role of CLASP1/2 in the maturation of the lytic IS in NK cells, and presents evidence supporting the contribution of GDMTs in this process.

**Summary sentence:** The Golgi apparatus (GA) functions as a microtubule-organizing center (MTOC) in NK cells. During the recognition of tumoral cells by NK cells, CLASP1/2-mediated stabilization of GA-derived microtubules (GDMTs) facilitates vesicular LFA-1 (LFA-1_v_) trafficking toward the interaction surface, thereby promoting the immune synapse (IS) maturation.

## INTRODUCTION

Natural killer (NK) cells are essential cytolytic effectors of the innate immune system. They serve as the first line of defense, playing a crucial role in identifying and eliminating virally infected and cancerous cells. NK cell cytotoxicity relies on the formation of a specialized junction with target cells, known as the immune synapse (IS). Unlike other lymphocytes, NK cell activation is primarily regulated by the balance of interactions between multiple activating and inhibitory surface receptors and their corresponding ligands on target cells, which occur at the IS. Persistent outside-in signaling, generated by the interaction of NK activating receptors with their ligands at the IS in the absence of inhibitory signals, initiates a multistep process. This process culminates in the translocation of the centrosome and lytic granules to the IS, followed by the directed secretion of lytic molecules from the granules into the target cell (Rak et al., 2011). The initial contact between NK and target cells may involve various receptors, among which the integrin leukocyte functional antigen (LFA)-1 is of particular significance. LFA-1, a heterodimer composed of the αL (CD11a) and β2 (CD18) chains, is essential for stabilizing the IS (Hoffmann et al., 2011). Moreover, LFA-1 contributes to the NK cytotoxic response by initiating signaling pathways that promote lytic granule convergence to the centrosome and the polarization of the lytic granules/centrosome towards the IS (Barber et al., 2004; Bryceson et al., 2005).

Downstream of the initial interaction between NK cell receptors and their specific ligands, the cytoskeleton plays a crucial role in NK cell cytolytic activity. The role of microtubules in NK-IS maturation has traditionally been associated with the delivery of lytic granules to the synaptic cleft (Trambas, C. M., & Griffiths, G. M., 2003). Our recent studies revealed an additional role for microtubules in NK cells, demonstrating that they participate in LFA-1 organization at the IS (Pariani et al., 2023). These studies further revealed the existence of an intracellular pool of LFA-1 that polarizes toward the IS in a Golgi-dependent manner, uncovering a Golgi/microtubule-dependent pathway that ensures LFA-1 organization at the IS and supports IS maturation.

Concomitantly with their central role in IS maturation, both the actin and the microtubule cytoskeleton suffer extensive remodeling during NK cell activation. While the factors governing actin cytoskeleton remodeling during NK activation have been thoroughly characterized, the regulation of microtubule dynamics has been relatively overlooked (Lagrue et al., 2013; Lee et al., 2021). Microtubules formation implies the polymerization of alpha/beta tubulin dimers. The dynamic instability associated with this process gives microtubules an intrinsic polarity, characterized by a highly dynamic, fast-growing plus end and a more stable minus end, which is often anchored to various cellular structures (Wu, J., & Akhmanova, A., 2017). The primary sites of microtubule minus end attachment are the microtubule organizing centers (MTOCs), which are responsible for microtubule nucleation. MTOCs are characterized by the presence of the γ-tubulin ring complex (γ-TuRC), a core component of the microtubule nucleation machinery that serves as a template for initiating microtubule polymerization. The best characterized MTOCs in animal cells are the centrosomes, which generate radial arrays of microtubules. In the last two decades, multiple studies in different cell types have characterized the role of the Golgi apparatus as a MTOC that generates asymmetric arrays of microtubules, which are better suited for contributing to cell polarity (Sanders and Kaverina, 2015). Whereas the centrosome is generally regarded as the primary MTOC in NK cells, the role of the Golgi apparatus as an MTOC in NK cells has not been assessed.

Microtubule nucleation at the Golgi requires the expression of A kinase anchoring protein 350 (AKAP350, AKAP450 or CG-NAP) at the cis-Golgi, which, along with myomegalin and/or CDK5Rap2 and CEP215, recruits γ-TuRC to the Golgi apparatus, facilitating microtubule nucleation (Rivero et al., 2009; Choi et al., 2010; Roubin et al., 2012; Sanders et al., 2017). The plus ends of GDMTs are stabilized at the trans-Golgi by the microtubule plus-tip interacting proteins CLASP1/2 (Efimov et al., 2007). CLASP1/2 were first isolated as proteins that associate with CLIP-170 and CLIP-115 at microtubule plus ends, which had microtubule-stabilizing properties on subsets of microtubules directed towards the leading edge in migrating fibroblasts (Akhmanova, 2001). Additional studies confirm those initial findings, and further demonstrate that CLASPs also localize to the centrosomes and the Golgi, where they increase the efficiency of microtubule nucleation (Efimov et al., 2007; Miller et al., 2009; Mimori-Kiyosue et al., 2005). While existing evidence supports the involvement of CLASP1/2 in stabilizing specific microtubule subsets microtubules involved in cell polarization (Lawrence and Rice, 2020), their role in the acquisition of cell polarity in lymphocytes has not been characterized.

Based on the background outlined above, this study aims to investigate the participation of CLASP1/2 in the maturation of NK-IS, and to evaluate the potential role of GDMTs in this process.

## MATERIALS AND METHODS

### NK and target cell lines

The immortalized NK YTS cells (Drexler and Matsuo, 2000) and KT86 cells, derived from the MHC class I-negative K562 erythroleukemia cell line stably expressing CD86 (Parry et al., 2003) were obtained from Dr. Jordan Orange lab (Texas Children’s Hospital, Houston, TX, USA). Cells were maintained in RPMI medium 1640 supplemented with 10% FBS, 1% L-glutamine, 1% non-essentials amino acids and 1% streptomycin/ampicillin mixture at 37°C and 5% CO_2_ atmosphere.

### *Ex vivo* NK cell purification

Blood samples from volunteer donors were used to prepare *ex vivo* NK cells. All human samples were obtained after informed patient consent and were used with approval of the institutional internal review board for the protection of human subjects of the Hospital Provincial del Centenario (Rosario, Argentina). In order to separate red blood cells from leukocytes, 20 ml of whole blood sample was centrifuged at 900 g for 15 minutes and the buffy coat was obtained. The buffy coat was resuspended in an equal volume of PBS and the Peripheral Blood Mononuclear Cells (PBMC) fraction was obtained by centrifugation on a Ficoll cushion for 30 minutes at 600g. The PBMC were washed twice with physiological solution and centrifuged for 15 minutes at 400g. The PBMC were resuspended in 500 uL of whole blood and incubated for 20 minutes at room temperature with RosetteSep Human NK cells Enrichment cocktail (StemCell # 15025) to purify NK cells by negative selection. After the incubation period, one volume of physiological solution supplemented with 2% of FBS was added and the NK cells enriched fraction was obtained by centrifugation on a Ficoll cushion for 20 minutes at 400g. Purified NK cells were washed twice in physiological solution supplemented with 2% FBS for 10 minutes at 400g, resuspended and maintained in RPMI medium 1640 supplemented with 10% FBS, 1% L-glutamine, 1% non-essentials amino acids, 1% streptomycin/ampicillin mixture and IL-2 500 UI/ml at 37°C and 5% CO_2_ atmosphere.

### Reduction of CLASP1 and CLASP2 expression by shRNA

In order to reduce both CLASP1 and CLASP2 expression in YTS and in *ex vivo* NK cells we used two different combinations of shRNA and proceeded as we described (Pariani et al., 2023). Briefly, constructs were made by annealing and ligating appropriate oligonucleotides containing CLASP1A or CLASP2 mRNA specific sequence or its scrambled control, into the AgeI and EcoRI cloning sites of pLKO.1-puro vector (details at http://www.addgene.org). These were sequenced and used to co-transfect human embryonic kidney 293 FT cells with Virapower lentiviral packaging mix (Invitrogen, Carlsbad, CA). Cells were allowed to produce virus for 24 hr. Media containing viruses were collected and used to directly transduce YTS or *ex vivo* NK cells overnight. The cells were allowed to recover for 24 h and then subjected to puromycin selection (2μg/ml) for 1 week. We obtained two different YTS cell populations with reduced expression of CLASP1 and CLASP2, YTS 1A2B and YTS 1B2A. The 1A2B cell line combined CLASP1A shRNA (5’-CCATTATGCCAACTATCT-3’) and CLASP2B shRNA (5’-GACATACATGGGTCTTAGA-3’), whereas 1B2A cell line combined CLASP1B shRNA (5’-CCTGAATTTACCATGTTACTT-3’) and CLASP2A shRNA (5’-GCCGTCTTGGTTGAAACTATT-3’), using the targeting sequences of siRNAs designed by Mimori-Kiyosue et al (2005). ex vivo NK cells were transduced with the viral particles for expression of the CLASP1B and CLASP2A targeting shRNAs. Silencing was confirmed by Western blot.

### Ice recovery assay

A total of 2×10^5^ YTS cells were placed on poly-l-lysine-coated coverslips and incubated for 30 minutes at 37°C. Then, coverslips were incubated on ice 50 min and allowed to recover at room temperature. For analysis of Golgi nucleation of microtubules, cells were recovered for 5 minutes and immediately treated 45 s with extraction buffer (60 mM PIPES, 25 mM HEPES, 10 mM EGTA, 2 mM MgCl2, 0.1% Tritón X-100, pH 6.9, supplement with 0.25 nM nocodazole and 0.25 nM paclitaxel). Cells were then fixed with methanol and stained with anti-GM130 or anti-Giantin (TA10, produced by the Institut Curie platform) and anti ɑ-tubulin antibody (Sigma-T9026). ɑ-tubulin association with the Golgi and the characterization of GDMTs were carried out as described in the ‘image analysis’ section.

### Immunofluorescence

KT86 cells were washed with PBS, resuspended at 2 ×10^6^ cells/ml and labeled with 300 nM Cell TrackerTM Deep Red (Invitrogen) for 15 min at 37°C. The reaction was stopped by the addition of an equal volume of fetal bovine serum (FCS), followed by a 2-min incubation at room temperature. Labeled target cells were washed twice and resuspended in RPMI complete medium. Conjugates between YTS cells and KT86 at a 2:1 ratio were established in suspension for 15 min at 37°C and adhered to poly-lysine-coated glass slides (Polyprep; Sigma-Aldrich) for additional 15 min. Non adherent cells were washed and cells adhering to the slide were fixed with 4% PFA at room temperature or methanol at −20°C. Fixed cells were blocked with 1% bovine serum albumin and 0.3%Triton X-100 in PBS, pH 7.4, for 10 min. Then, they were incubated for 2 h with mouse monoclonal antibody anti-LFA-1 (Biolegend 301202), anti-CLASP1 (Abcam, ab108620), anti-Perforin (Santa Cruz SC-136994), anti-GM130 (Abcam, ab52649) or anti-α tubulin (Sigma T5168) and rabbit monoclonal antibody anti γ-tubulin (Sigma-T5192). The coverslips were washed, incubated for 1 h with the secondary antibodies conjugated to Alexa 488, Alexa 560 or Alexa 633 (Molecular probes-A34055, 1:200) and with 4′,6-diamidino-2-phenylindole (DAPI) and mounted with ProLong (Invitrogen). Fluorescence was detected using LSM880 confocal with an ObserverZ1 inverted microscope. Serial optical 0.4 µm thick sections were collected in the z-axis. Z-stacks were built, and projections were obtained using ImageJ tools. To improve visualization of microtubule nucleation at the Golgi apparatus the open-source plug-in of imageJ, Super-Resolution Radial Fluctuation (SRRF), was used for the α-tubulin channel to enhance microtubule image resolution (Gustafsson *et al.,* 2016). First, for the acquisition of the images, the *x,y* plane where the Golgi signal was maximum was chosen and a time series tool was applied to obtain a sequence of 100 image frames with an interval of 2.53 seconds. Finally, the α-tubulin channel was chosen and analyzed by the SRRF plug-in. In preparing the figures, adjustment in brightness and contrast were equally applied to the entire images using Adobe Photoshop software to improve visualization of fluorescence.

### Image analysis

Lytic granule convergence and centrosome and lytic granule polarization towards the IS. YTS:KT86 conjugates were permeabilized and stained with anti γ-tubulin, perforin and DAPI, for lytic granules, centrosome and nucleus identification, respectively. The IS was defined as the cell to cell contact region identified in the images obtained by differential interference microscopy (DIC). The cell perimeter of the YTS cell forming the IS was drawn using the same images. At the *x,y* plane corresponding to the centrosome center, a threshold was set on the perforin or γ-tubulin channel to define the respective masks, which were used to automatically outline the centrosome and the lytic granule regions. The *x,y* coordinates values for the IS, the lytic granules, the centrosome, and the cell centroids were determined using the appropriate ImageJ tool and the distances from the lytic granules, the centrosome or the cell centroids to the IS centroid, or from the lytic granules to the centrosome centroid were calculated. In the case of lytic granules, an average area-weighted distance (AWD) was calculated for each cell by applying a modified use of Shepard’s Method:

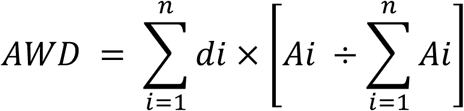

where *Ai* is the area of each particular lytic granule region and *di* is the distance from this particle to the IS or the centrosome. This modification allowed weighting the distances by the area of each lytic granule region, which is important considering that, when they are close together, clusters of lytic granules could inappropriately be discerned as an individual granule of a larger area and, therefore, if a factor considering each granule area is not used, those granules could be under-represented. At least 30 cells from three independent experiments were analyzed for each condition.

Polarization of LFA-1-vesicles towards the IS. YTS:KT86 or *ex vivo* NK:KT86 cell conjugates were permeabilized and stained with anti LFA-1, for identification of LFA-1vesicles. The IS and the cell regions were defined in the DIC images, as explained above. A threshold was set on the LFA-1 channel to define a mask, which was used to automatically outline regions corresponding to the LFA-1 vesicles. The IS, LFA-1 vesicles and the cell centroid were determined using the appropriate ImageJ tool and the distances from the LFA-1 vesicles or the cell centroids to the IS centroid were calculated. Similar to what we explained above for lytic granules, an average area-weighted distance (AWD) was calculated for each cell.

LFA-1 accumulation at the IS. YTS or *ex vivo* NK cells conjugated with KT86 cells were analyzed. DIC images were used for IS and cell selections as we describe above. For Fig. 4a, the area of the LFA-1 positive region was additionally determined in the LFA-1 channel. Total intensity of fluorescence was measured at NK-IS and at the whole cell and the percentage of protein accumulation at the IS was calculated.

For estimating α-tubulin association to the Golgi apparatus, a threshold on GM130 channel or Giantin was used to define a mask, which was utilized to automatically outline the Golgi apparatus region. The cell region was delimited in the DIC channel. Total intensity of fluorescence in the α-tubulin channel was measured in each region of interest, and the percentage of α-tubulin fluorescence at the Golgi apparatus was calculated for each cell. The number and length of microtubules were counted and measured manually in images obtained by confocal microscopy. The (x,z) reconstructions of the images at y levels where the Golgi did not overlap with the centrosome, were obtained using the Reslice tool of ImageJ for better visualization. Microtubules were identified in the α-tubulin channel, and GDMTs were recognized by their partial colocalization with Golgi fragments. The length of microtubules was measured using the segmented line tool from ImageJ.

### Immunoblotting

Cells were harvested at 400 *g* for 5 min at room temperature and washed with cold PBS. Pelleted cells were lysed in ice-cold lysis buffer (50 mM Tris-HCl [pH 7.5], 100 mM NaCl, 15 mM EDTA and 1% Triton X-100, with protease inhibitors) and subjected to two freeze– thaw cycles. Lysates were centrifuged at 1000 *g* for 5 min at 4°C, and the clear supernatants were conserved.

For all samples, total protein concentrations were measured according to Lowry *et al*. (1951). Samples were heated for 10 min at 90°C in sample buffer (20 mM Tris-HCl, pH 8.5, 1% SDS, 400 μM DTT, 10% glycerol). Samples containing equal amounts of proteins were subjected to SDS polyacrylamide gel electrophoresis. The proteins were transferred to nitrocellulose membranes (Amersham Pharmacia Biotech). Blots were blocked with 5% non-fat dry milk in PBS, 0.3% Tween-20 (PBS-Tween). Nitrocellulose blots were then probed with the monoclonal mouse anti-CLASP1 antibody (Abcam, ab108620), rat anti-CLASP2 antibody (Abcam, ab95373), mouse anti-AKA350 (14G2, Schmidt et al 2001) and mouse monoclonal antibodies anti-α-tubulin (Sigma-T5168), anti-β actin (Sigma A2228), anti-GAPDH (sc-47724) or anti-CIP4 (BD Bioscience-612556) as loading controls. The blots were washed and incubated with the horseradish peroxidase-conjugated corresponding secondary antibodies, and bands were detected by enhanced chemiluminescence (Pierce, Thermo Scientific). Blot images were obtained using Amersham ImageQuant500 (Cytiva). The bands were quantitated by densitometry using the NIH Image J program.

### Statistical analysis

Data are expressed as mean ± s.e.m. Paired Student’s t-test or nonparametric Mann–Whitney test were used for comparison between groups when necessary. For multiple comparisons, one-way ANOVA followed by Tukey’s multiple comparisons test was used. p < 0.05 was considered statistically significant.

## RESULTS

### 1. CLASP1/2 participate in lytic IS maturation

We generated two YTS cell lines with decreased expresión of CLASP1/2 (CLASPKD) expressing different combinations of shRNAs (1A2B and 1B2A) targeted to specific sequences of CLASP1 and CLASP2, which have been demonstrated to effectively knock down CLASP1 and CLASP2 expression (Mimori-Kiyosue *et al*., 2005). Western blot analysis of cell lysates showed that the expression of both CLASP1 and CLASP2 proteins was significantly lower in the YTS CLASPKD 1B2A cell line (94% and 82% reduction, respectively; Figure 1a), whereas in the YTS CLASPKD 1A2B cell line, CLASP knockdown was only effective for CLASP1 (−62%). To analyze the participation of CLASP1/2 in the NK cell effector response against tumor cells, we examined the maturation of the immunological synapse (IS) between control or CLASPKD YTS cells and erythroleukemia-derived KT86 cells. The convergence of lytic granules toward the centrosome and the polarization of the centrosome and lytic granules to the NK-target cell interaction zone during IS formation are key events in the NK cytolytic response. Immunofluorescence confocal microscopy analysis of YTS-KT86 cell conjugates (Figure 1b) showed that the distance from the centrosome or the lytic granules to the IS was increased in CLASP1/2 KD cells (Figures 1c and 1d). Concurrently, the most significant inhibition in centrosome and lytic granule polarization corresponded to the CLASPKD 1B2A YTS cell line, reflecting the efficiency of CLASP1/2 knockdown. On the other hand, lytic granule convergence to the centrosome was not impaired in YTS cell lines with decreased expression of CLASPs (Figure 1e). Additionally, ex vivo cultures of freshly isolated NK cells were transduced with lentiviral particles for the expression of CLASP1B and CLASP2A shRNAs. Analysis of protein levels by western blot indicated that the expression of both CLASP1 and CLASP2 was decreased (−90% and −40%, respectively; Figure 1f). Examination of ex vivo NK-KT86 cell conjugates revealed that the centrosome polarized to the immunological synapse (IS) in control cells but not in CLASPKD NK cells (Figure 1g,h). Furthermore, the polarization of lytic granules to the IS was impaired in ex vivo NK CLASPKD cells (Figure 1i). Consistent with observations in YTS cells, the convergence of lytic granules to the centrosome in ex vivo NK cells was not affected by the reduction in CLASP1/2 expression (Figure 1j). These results indicate that CLASP1/2 are necessary for the maturation of the lytic IS, participating in events that have impact downstream of lytic granule convergence to the centrosome and upstream of centrosome polarization to the IS.

**Figure 1.**
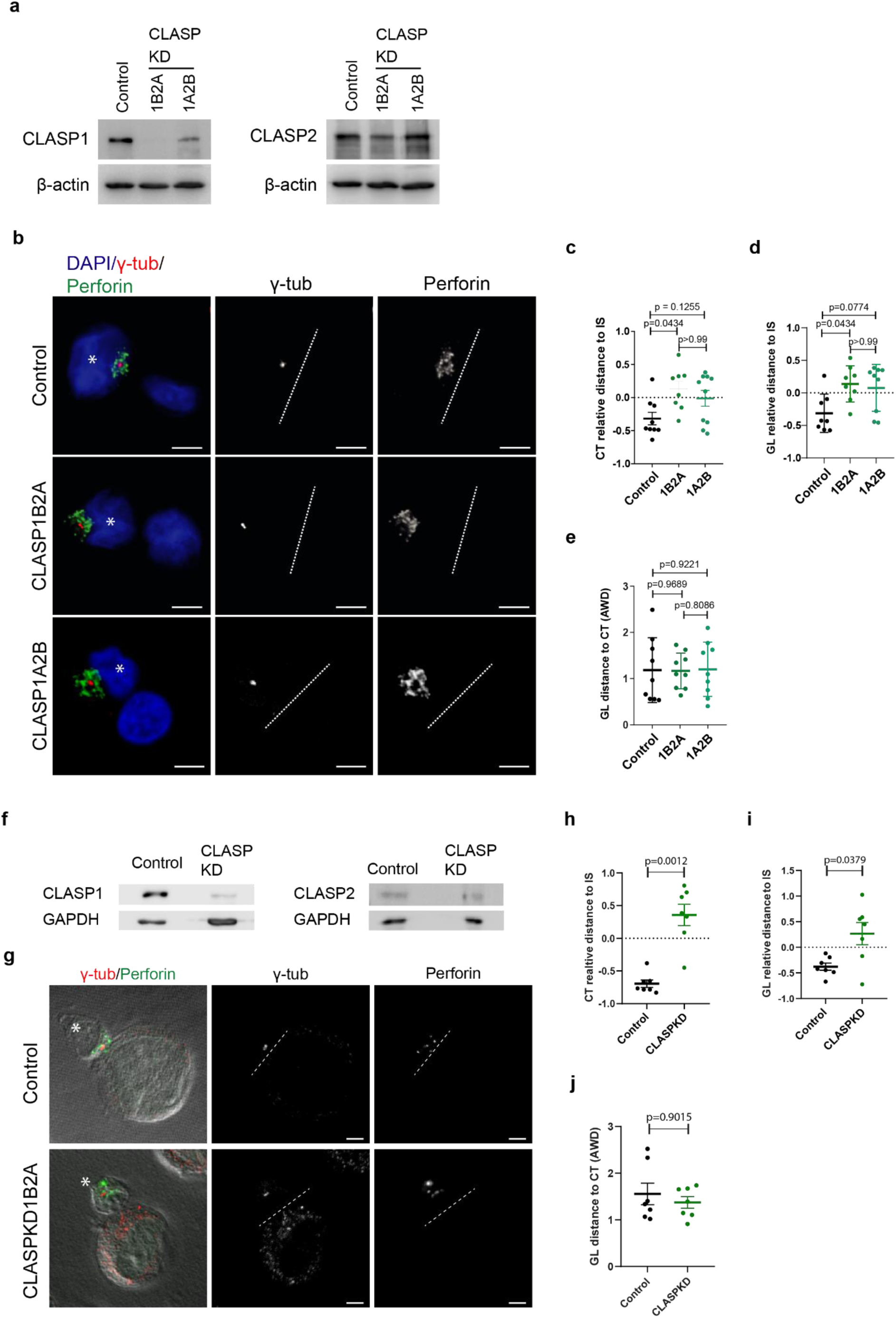
Reduction in CLASP1/2 expression inhibits IS maturation. (a-e) Two different combinations of shRNAs for CLASP1 and CLASP2 were used to generate two YTS cell lines with decreased expression of CLASP1 and CLASP2 proteins (1A2B and 1B2A) as described in the Material and Methods section. a) Western blot analysis of CLASP1 and CLASP2 expression in control and 1A2B and 1B2A cell lines. β-actin was used as loading control. (b-e) Control or CLASPKD YTS cells were co-cultured with KT86 target cells for 30 minutes. After this period, cells were fixed, stained and analyzed by confocal microscopy. b) Merge images show staining for perforin in green, γ-tubulin in red and DAPI in blue. YTS cells are marked with an asterisk. Individual channels for perforin and γ-tubulin are shown in greyscale. Dashed lines mark the IS surface. Scale bars, 5 μm. c-e) Dot plots represent the relative distance of centrosome to the IS for each cell (c) the mean AWD of lytic granules (GL) to the IS (d) and the mean AWD from the LG to the centrosome (e) for each cell for one experiment, representative of 3 experiments. (f-j) ex vivo NK cells with reduced expression of CLASP1/2 proteins were prepared as described in the Materials and Methods section. f) Western blot analysis of CLASP1 and CLASP2 expression in control and in 1B2A ex vivo NK cells. GAPDH was used as loading control. (g-j) Control CLASPKD or ex vivo NK cells were co-cultured with KT86 target cells for 30 minutes. The conjugates were fixed, stained and analyzed by confocal microscopy. g) Merge images show Perforin in green, γ-tubulin in red and DIC in greyscale channels. NK cells are marked with an asterisk. Individual channels for Perforin and γ-tubulin are shown in greyscale. Dashed lines mark the IS surface. Scale bar, 2.5 μm. h-j) Dot plots represent the relative distance of centrosome to the IS for each cell (h) the mean AWD of lytic granules (GL) to the IS (i) and the mean AWD from the LG to the centrosome (j) for each cell for one experiment, representative of 3 experiments. Error bar represents SEM.

### 1. CLASP1/2 are necessary for LFA-1 accumulation at the IS

LFA-1 reorganization at the immunological synapse (IS) is an essential event in NK cell activation, which can lead to centrosome polarization. We have previously identified a mechanism of LFA-1 relocalization towards the IS during NK activation that requires the integrity of microtubules and the Golgi apparatus (Pariani *et al*., 2023). To elucidate whether CLASP1/2 are involved in this mechanism, we further analyzed LFA-1 localization during CLASP1KD YTS cells recognition of tumor cells. Analysis of intracellular LFA-1 by immunofluorescence confocal microscopy showed impaired polarization of LFA-1 vesicles toward the IS edge in CLASPKD cells (Figure 2a, b). Concurrently, the accumulation of LFA-1 at the IS was inhibited in both CLASPKD cell lines (Figure 2a, c). Consistent with the efficiency of CLASP1/2 knockdown, the most significant effect on LFA-1 reorganization was observed in the CLASPKD 1B2A cell line (Figure 2b, c). These results indicate that CLASPs are essential for LFA-1 reorganization toward the IS during NK cell activation.

**Figure 2.**
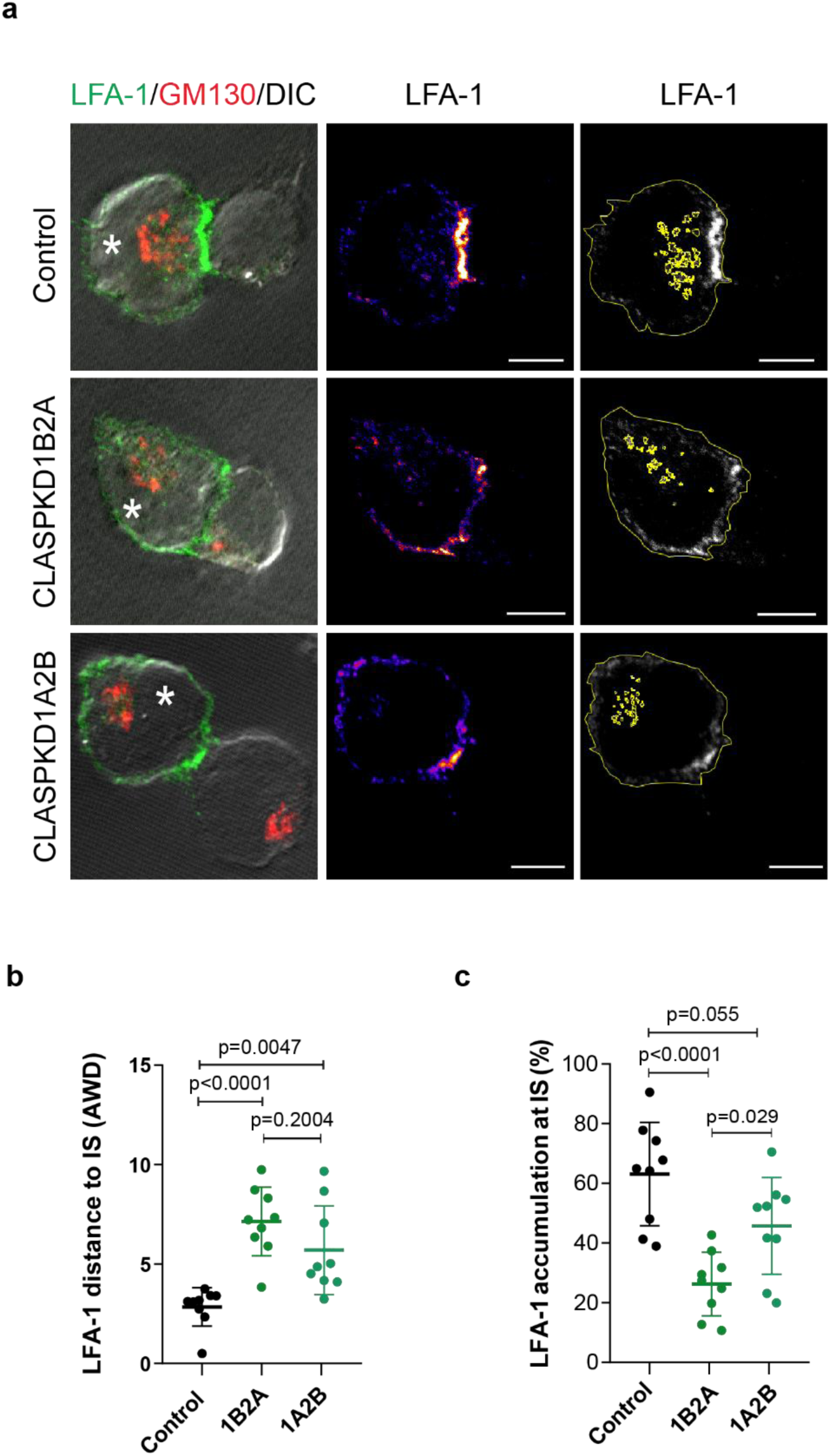
CLASPs1/2 participate in LFA-1 organization at the IS. Control or CLASPKD YTS cells were co-cultured with KT86 target cells for 30 minutes. The conjugates were fixed, stained and analyzed by confocal microscopy. Merge images show LFA-1 in green, Giantin in red and DIC in grayscale. LFA-1 accumulation is shown in pseudocolor. LFA-1 channel is shown in grayscale with the ROIs delimiting the cell and individual vesicles in yellow. The asterisk marks YTS cells. (b) The dot plot represents the mean distance (AWD) of LFA-1 vesicles to the IS for each cell in one experiment, representative of three experiments. (c) The dot plot represents the percentage of f LFA-1 fluorescence that localizes at the IS for each cell in one experiment, representative of 3 experiments. Error bars represent SEM. Scale bar, 5 μm.

### 2. Microtubule nucleation at the Golgi Apparatus

Similarly to our findings on CLASP1/2 involvement in NK-IS maturation, our previous studies demonstrate that AKAP350 participates in LFA-1 reorganization at the NK-IS, facilitating NK cytolytic function (Pariani et al, 2023). Considering that, as well as CLASP1/2, AKAP350 have a central role in the nucleation of GDMTs, we aimed to evaluate the putative function of the Golgi as a MTOC in NK cells, which has not been previously reported. The immunofluorescence analysis of ice recovery assays revealed the presence of newly nucleated microtubules at the Golgi, whose arrays were clearly different from the radial arrays of centrosome-nucleated microtubules, in YTS cells in the early recovery phase (Figure 3a). Similarly, *ex vivo* NK cells showed α-tubulin association with the Golgi at 3 minutes of recovery, confirming the GA’s capacity to nucleate microtubules in NK cells (Figure 3b). Additionally, CLASP1 staining of ex vivo NK cells demonstrated colocalization with α-tubulin at the Golgi apparatus (Figure 3c), which is consistent with a role of CLASP1 in stabilizing GDMTs in NK cells.

**Figure 3.**
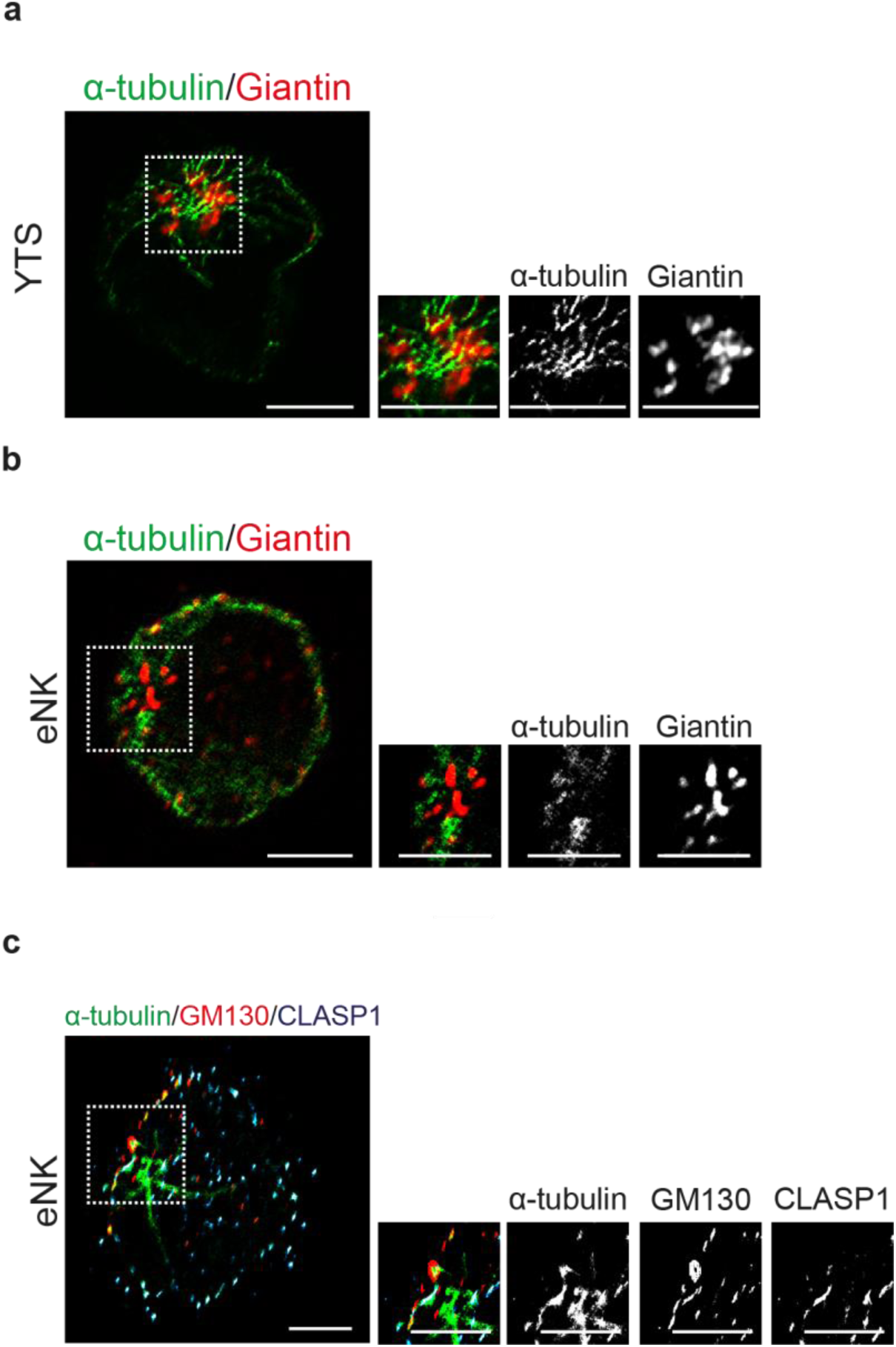
Microtubule nucleation at the GA. YTS (a) or *ex vivo* (b-c) NK cells were seeded on coverslips coated with poly-l-lysine for 30 minutes and then subjected to an ice recovery assay as described in materials and methods. Merge images of YTS (a) or *ex vivo* (b) NK cells show the staining of α-tubulin (green) and the Golgi marker Giantin (red). The inset shows merge and individual channels of a higher amplification of the boxed area. Individual channels are shown in grayscale .c) Merge images of *ex vivo* NK cells show the staining of α-tubulin (green), CLASP1 (blue) and the Golgi marker GM130 (red). The inset shows merge and individual channels of a higher amplification of the boxed area. Individual channels are shown in grayscale. Scale bars, 5 μm in **a** and 2.5 μm in **b**.

### 3. CLASP1/2 and AKAP350 participate in microtubule nucleation at the Golgi apparatus in NK cells

In epithelial cells, CLASP1 and CLASP2 specifically participate in the stabilization of microtubules at the Golgi apparatus (Efimov et al., 2007). To investigate the role of CLASP1/2 in microtubule nucleation at the Golgi in NK cells, we conducted ice recovery assays using both control and CLASPKD YTS cells. Confocal microscopy analysis revealed that microtubules in CLASPKD cells exhibited a significantly reduced number of GDMTs compared to control cells (Figure 4a,b). Furthermore, GDMTs in CLASPKD cells were notably shorter in CLASPKD cells (Figure 4c). Correlating with the efficiency of CLASP1 and CLASP2 knockdown, the most significant decrease in GDMTs number and length was observed in the YTS 1B2A cell line. These results indicate that, in accordance with CLASP1/2 function in other cell types, in NK cells GDMTs are stabilized by CLASP1/2 early during their nucleation.

**Figure 4.**
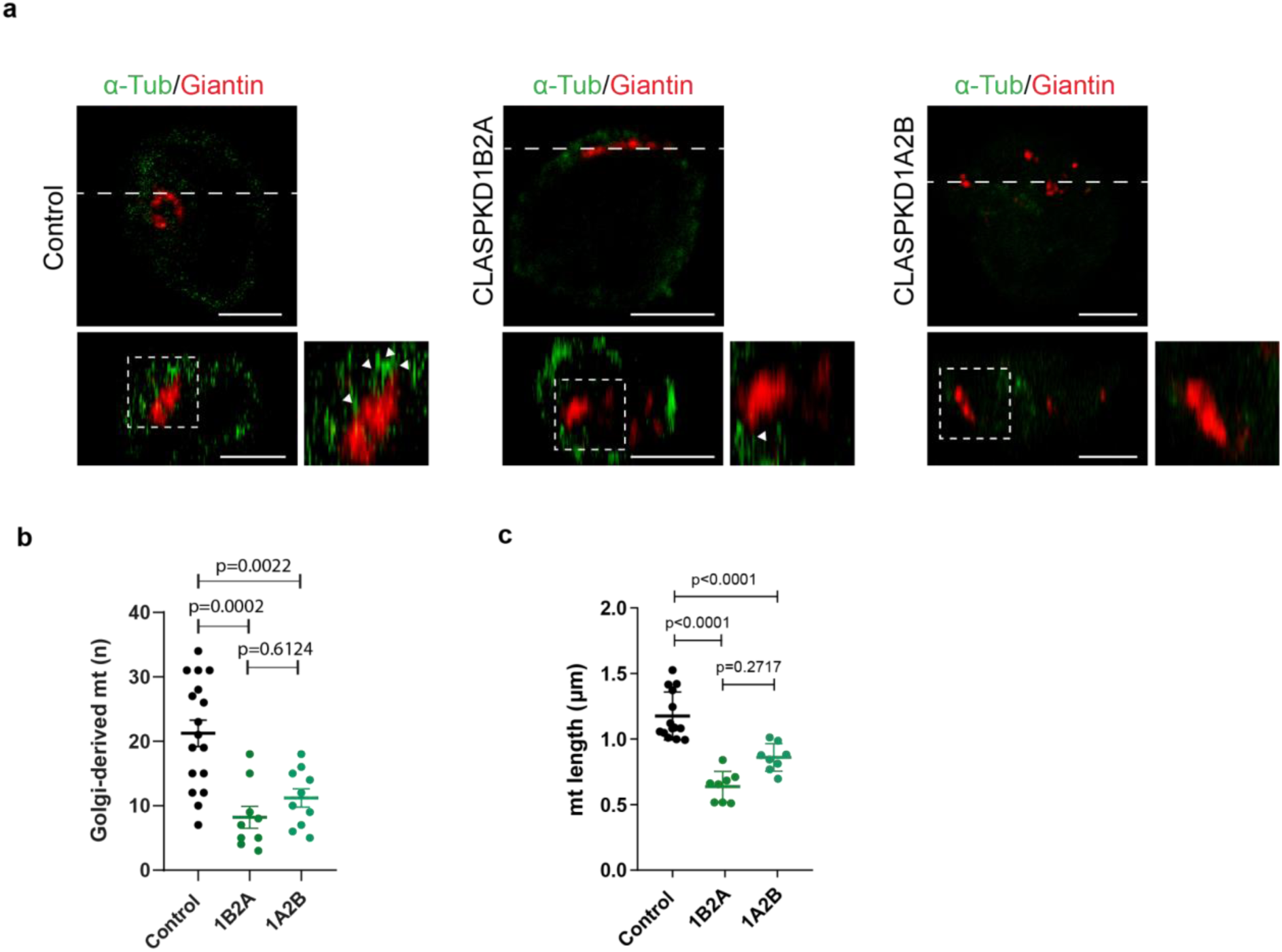
Reduction of CLASP1/2 expression inhibits microtubule nucleation at the Golgi. Control or CLASPKD YTS cells were incubated on coverslips coated with poli-L-lysine for 30 minutes at 37 °C and then subjected to ice recovery assays, fixed and stained. a) Merge images show giantin (red) and α-tubulin (green) staining for representative *x,y* planes (above) and for the orthogonal views of the x,z reslices (below) at the ordinate level that is indicated with dashed lines in the *x,y* plane, which were used for better quantification of GDMTs. The insets show amplifications of the boxed areas of the orthogonal views. Arrowheads indicate GDMTs. (b,c) Dot plots represent the number of microtubules (b) and the mean length of microtubules (c) originated from the Golgi for each cell in one experiment, representative of 3 experiments. Error bars represent SEM. Scale bar, 5 μm.

AKAP350 is a centrosome/Golgi apparatus associated scaffold protein that participates in microtubule dynamics (Larocca *et al.,* 2006; Almada *et al*., 2017). In epithelial cells, AKAP350 has a central role in the nucleation of microtubules at the Golgi apparatus (Rivero *et al.,* 2009). We generated YTS cells with decreased expression of AKAP350 using a lentiviral short hairpin RNA (shRNA) expression system, as previously described (Pariani et al, 2023). The reduction in AKAP350 expression was verified by western blot (Figure 5a). Analysis of microtubule distribution indicated a reduction in microtubule association with the Golgi in AKAP350 knockdown (AKAP350KD) cells (Figure 5b). Moreover, ice recovery assays showed that both the α-tubulin association with the Golgi and the number of microtubules originating from this organelle were decreased in AKAP350KD cells, indicating that the reduction in AKAP350 expression impaired microtubule nucleation at the Golgi (Figure 5c). On the other hand, neither the fraction of α-tubulin associated with the centrosome nor the number of microtubules nucleated at this organelle were affected by AKAP350 knockdown. Therefore, similar to what has been demonstrated in epithelial cells, the reduction in AKAP350 expression in NK cells specifically inhibits microtubule nucleation at the Golgi without significantly impacting centrosomal microtubule nucleation. Altogether, our data show that AKAP350 and CLASPs are both necessary for formation of GDMTs in NK cells. Because this is the only known common function for AKAP350 and CLASPs, we conclude that IS polarization defects observed upon depletions of either of these proteins (this study and Pariani et al, 2023) result from the reduced GDMT numbers and that GDMTs are essential for proper IS formation.

**Figure 5.**
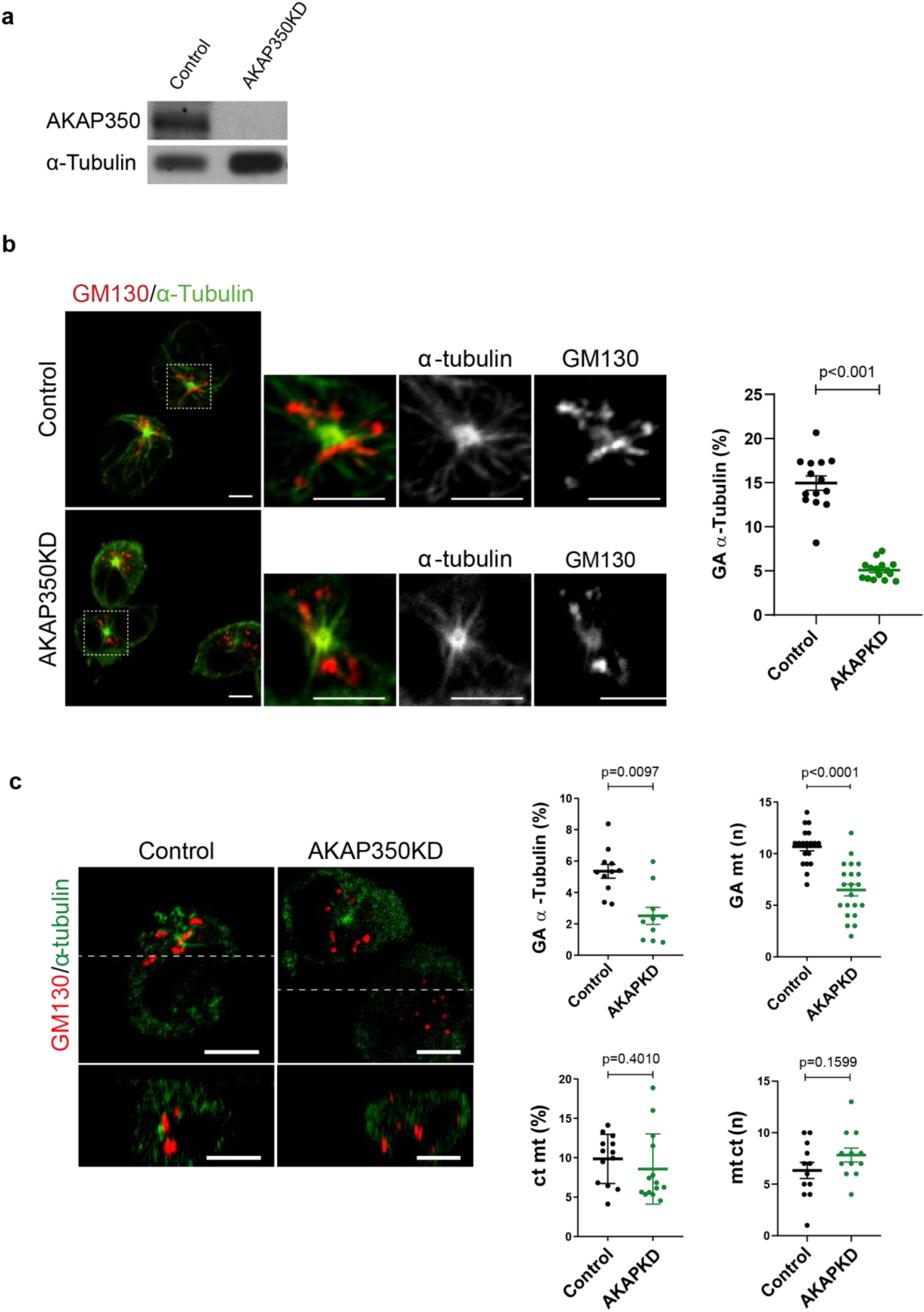
AKAP350 participates in microtubule nucleation at the GA. YTS Cells with decreased expression of AKAP350 were prepared by using a lentiviral system for shRNA expression as described in the Materials and Methods section. a) Western blot analysis of AKAP350 expression in control or AKAP350KD YTS cell lines. CIP4 was used as loading control. (b-c) Control or AKAPKD YTS cells were incubated on coverslips coated with poli-L-lysine y for 30 minutes and then fixed (b) or subjected to ice recovery assays and fixed at 3 min of recovery (c). Merge images show staining for GM130 (red) and α-tubulin (green). In (c), images also show the orthogonal views of the x,z reslice at the ordinate level that is indicated with dashed lines, which were used for better quantification of GDMTs. Dot plots represent the mean fraction of α-tubulin fluorescence associated to the Golgi apparatus (b,c) and to the centrosome (c), expressed as percentage of total cell fluorescence for the α-tubulin channel (b,c), and the mean number of microtubules derived from the centrosome or from the Golgi apparatus for each cell (c), in one experiment, representative of 3 experiments. At least 10 cells were analyzed in each experiment. Error bars represent SEM. Scale bars, 5 μm.

## DISCUSSION

CLASP1/2 belong to a group of proteins that bind to microtubule plus ends, regulating their dynamic stability. More specifically, CLASP1/2 bind to specific networks of microtubules, such as GDMTs, enhancing their stability and thereby ensuring the establishment of cell polarity in migrating cells and neurons (Lawrence and Rice, 2020). Centrosome polarization is a critical checkpoint during the acquisition of IS-cell polarity during NK cytolytic response: if signals from inhibitory NK receptors surpass those from activating receptors before centrosome translocation, the IS formation ceases (Hsu et al., 2016). Conversely, if NK activating receptor signals prevail, the centrosome and lytic granules move toward the IS, facilitating granule secretion into the synaptic cleft. LFA-1 outside-in signaling alone is sufficient to stimulate lytic granule convergence, centrosome polarization, and directed secretion of the lytic granules (Barber et al., 2004; Bryceson et al., 2005). Notably, blockade of LFA-1 or insufficient ligation with ICAM-1 disrupts early signaling events, resulting in irregular IS formation and impaired NK cell cytotoxicity (Raemer et al., 2009; Barber et al., 2004; Eitler et al., 2024). Our data demonstrate that normal CLASP1/2 expression is crucial for LFA-1 reorganization at the NK-target cell interaction site, and for the efficient polarization of the centrosome and lytic granules toward the IS. Therefore, the current study presents novel findings that underscore the role of CLASP1/2 in the development of IS-cell polarity during NK cytolytic response against tumoral target cells.

In several cell types, the GDMTs are of utmost importance for polarized traffic of proteins and to maintain the cell polarity state (Guo *et al*., 2014; Yadav, S., & Linstedt, 2011; Zhu, X., & Kaverina, I., 2013). Therefore, we hypothesize that the participation of CLASP1/2 in LFA1 reorganization towards the NK-IS could be related to a role of CLASP1/2 in GDMTs stabilization. In immune cells, the centrosome has traditionally been regarded as the exclusive MTOC. Even though in migratory T cells, *Ong et al.* identified the presence of non centrosomal MTOCs that rely on the protein AKAP350 for microtubule nucleation, they could not identify the Golgi apparatus as one of those MTOCs (Ong *et al*., 2018). In this line, there are no previous studies that have reported a potential role of the Golgi as an MTOC in NK cells. Furthermore, the role of the Golgi in IS formation has long remained uncharacterized. According to our previous studies, the disruption of the microtubule cytoskeleton or the impairment of the Golgi function inhibited the polarized traffic of the intracellular pool of LFA-1 to the IS in NK cells (Pariani *et al*., 2023). Moreover, in those studies we demonstrated that both the decrease in AKAP350 expression and the displacement of AKAP350 from the Golgi impaired LFA-1 polarization towards the IS (Pariani *et al*., 2023). Nevertheless, we did not have direct evidence of the existence of a pool of GDMTs. In the current work, ice recovery assays in NK cells demonstrated the presence of newly nucleated microtubules at the Golgi, which were decorated by CLASP1. These data provide the first evidence of the Golgi functioning as a MTOC in NK cells, and support a model where GDMTs participate in LFA-1 traffic towards the IS, thus regulating IS maturation. As commented above, in epithelial derived cells CLASP1/2 stabilize GDMTs at the trans-Golgi. CLASP1/2 mediated stabilization of GDMTs ensures microtubule elongation and leads to the formation of the asymmetrical array of microtubules associated to this organelle, clearly distinct from the radial array featured by centrosomal microtubules (Vinogradova *et al*, 2009; Efimov *et al*., 2007). Our results showed that, in NK cells with decreased expression of CLASP1/2, microtubule nucleation a the Golgi was less efficient, and that GDMTs were shorter, which suggest that, in NK cells, CLASP1/2 are responsible for a stabilization mechanism similar to those described in other cell types. Several studies have reported that AKAP350 participates in microtubule nucleation at the Golgi in different experimental systems and organisms (Rivero *et al*, 2009, Ori-McKenney *et al*, 2012; Maia *et al*, 2013, Tonucci *et al*, 2018), which is linked to AKAP350 ability to recruit γ-TuRC either directly, or indirectly via CDK5Rap2 or its paralog myomegalin (reviewed in Sanders and Kaverina, 2015). In the current study, we found that, in NK cells, the decrease in AKAP350 expression reduced the number of microtubules emerging from the GA, and decreased α-tubulin colocalization with this organelle. Those results indicate that AKAP350 facilitates microtubule nucleation at the Golgi in NK cells. Overall, those results indicate that the Golgi functions as a MTOC in NK cells, through a nucleation process that shares key components with the Golgi nucleation of microtubules characterized in other cell types.

We previously demonstrated that reduced AKAP350 expression in NK cells disrupts centrosome and lytic granule translocation toward the IS, resulting in diminished cytotoxic activity (Pariani *et al,* 2023). The inhibitory effect of the decrease of AKAP350 on IS maturation was, at least partially, secondary to the defective reorganization of LFA-1 at the IS membrane in NK cells. Those studies also revealed that the initial steps in the IS formation, that is NK cell adhesion to target cells and the initial activation-signaling, were not affected by the decrease in AKAP350 expression, but that 30 min after the initial interaction, the activation signaling was significantly diminished in AKAP350KD cells. In the present study we found that depletion of CLASP1/2 did not affect lytic granule convergence towards the centrosome, implicating that these proteins are not involved in the initial phase of NK-IS establishment. Altogether, our results are consistent with a model in which GDMTs facilitate the transport of LFA-1 to the IS, where, through interaction with its ligand on target cells, it generates activation signals that promote IS maturation, enabling the lytic response against the target cell. According to our previous studies and the hereby presented data, GDMTs-dependent LFA-1 reorganization at the IS is not required for NK cell adhesion to target cells or for the initial signaling that activates lytic granule convergence to the centrosome, but it is essential for the later polarized enrichment of LFA-1 at the IS surface, which strengthens the activation signaling that drives the final maturation of the IS.

In conclusion, this manuscript presents data that demonstrate that CLASP1/2 participate in the maturation of the lytic IS in NK cells and reveal that the Golgi functions as a MTOC in NK cells. Furthermore, this study provides evidence supporting a model where, during NK recognition of tumoral cells, GDMTs facilitate LFA-1 trafficking towards the SI thus enhancing activating signaling that leads to IS maturation and directed lytic granule secretion into tumoral cells.

## ACKNOWLEDGEMENTS AND FUNDING

This work was supported by Grants PIP2720 from CONICET, PICT2020-02817 from ANPCyT (to MCL) and National Institutes of Health (NIH) grants NIDDK R01DK106228 and NIGMS MIRA R35GM127098 (to IK). The funders had no role in study design, data collection and analysis, decision to publish, or paper preparation.

## AUTHOR CONTRIBUTIONS

All the experiments were performed at IFISE (CONICET-UNR), and at the Laboratory of biology and biochemistry of T. cruzi from the IBR (CONICET-UNR)

APP: conceptualization, investigation, analysis of the data, writing-original draft and revision of the manuscript.

VH, TRM, VA, EA, RV: investigation, writing-revision of the manuscript.

CF, JRG: conceptualization, writing-revision of the manuscript.

IK: conceptualization, writing-review, editing, revision of the manuscript.

MCL: conceptualization, analysis of the data, writing-original draft, review, editing and revision of the manuscript.

All the authors give their consent for publication.

## REFERENCES

Akhmanova A. CLASPs are CLIP-115 and-170 associating proteins involved in the regional regulation of microtubule dynamics in motile fibroblasts. Cell 2001:104:923–935. doi: 10.1016/s0092-8674(01)00288-4

Al-Bassam J, Kim H, Brouhard G, van Oijen A, Harrison SC, & Chang F. CLASP promotes microtubule rescue by recruiting tubulin dimers to the microtubule. Dev cell 2010:19:245–258. doi: 10.1016/j.devcel.2010.07.016.

Almada E, Tonucci FM, Hidalgo F, Ferretti A, Ibarra S, Pariani A, Vena R, Favre C, Girardini J, Kierbel A, Larocca MC. AKAP350 recruits EB1 to the spindle poles, ensuring proper spindle orientation and lumen formation in 3d epithelial cell cultures. Sci Rep 2017:7:1–15. doi: 10.1038/s41598-017-14241-y.

Barber DF, Faure M, Long EO. LFA-1 contributes an early signal for NK cell cytotoxicity. J Immunol 2004:173:3653–3659. doi: 10.4049/jimmunol.173.6.3653.

Bryceson YT, March ME, Barber DF, Ljunggren HG, Long EO. Cytolytic granule polarization and degranulation controlled by different receptors in resting NK cells. J Exp Med. 2005:202:1001–1012. doi: 10.1084/jem.20051143.

Carpier JM, Zucchetti AE, Bataille L, Dogniaux S, Shafaq-Zadah M, Bardin S, Lucchino M, Maurin M, Joannas LD, Magalhaes JG, Johannes L, Galli T, Goud B, Hivroz C.. Rab6-dependent retrograde traffic of LAT controls immune synapse formation and T cell activation. J Exp Med.:2018:215:-1265. doi: 10.1084/jem.20162042.

Choi YK, Liu P, Sze SK, Dai C, Qi RZ. CDK5RAP2 stimulates microtubule nucleation by the γ-tubulin ring complex. J Cell Biol. 2010:191:1089–1095. doi: 10.1083/jcb.201007030.

Drexler HG, Y Matsuo. Malignant hematopoietic cell lines: in vitro models for the study of natural killer cell leukemia-lymphoma. Leukemia 2000:14:777–782. doi: 10.1038/sj.leu.2401778

Efimov A, Kharitonov A, Efimova N, Loncarek J, Miller PM, Andreyeva N, Kaverina I. Asymmetric CLASP-dependent nucleation of noncentrosomal microtubules at the trans-Golgi network. Dev Cell 2007:12: 917–930. doi: 10.1016/j.devcel.2007.04.002.

Eitler J, Rackwitz W, Wotschel N, Gudipati V, Shankar NM, Sidorenkova A, & Tonn T. CAR-mediated targeting of NK cells overcomes tumor immune escape caused by ICAM-1 downregulation. J Immunother Cancer 2024:12. doi: 10.1136/jitc-2023-008155.

Guo Y, Sirkis DW, Schekman R. Protein sorting at the trans-Golgi network. Annu Rev Cell Dev Biol 2014:30:169–206. doi: 10.1146/annurev-cellbio-100913-013012.

Hoffmann SC, Cohnen A, Ludwig T, Watzl C. 2B4 engagement mediates rapid LFA-1 and actin-dependent NK cell adhesion to tumor cells as measured by single cell force spectroscopy. J Immunol 2011:186:2757–2764. doi: 10.4049/jimmunol.1002867.

Hsu HT, Mace EM, Carisey AF, Viswanath DI, Christakou AE, Wiklund M, … & Orange JS. NK cells converge lytic granules to promote cytotoxicity and prevent bystander killing. J Cell Biol. 2016:215:875–889. doi: 10.1083/jcb.201604136.

Larocca MC, Jin M, Goldenring JR. AKAP350 modulates microtubule dynamics. E J Cell Biol 2006:85:611–619. doi: 10.1016/j.ejcb.2005.10.008

Lagrue K, Carisey A, Oszmiana A, Kennedy PR, Williamson DJ, Cartwright A, Barthen C, Davis DM. The central role of the cytoskeleton in mechanisms and functions of the NK cell immune synapse. Immunol Rev 2013:256:203–221. doi: 10.1111/imr.12107

Lawrence EJ, Zanic M, Rice LM. CLASPs at a glance. J Cell Sci. 2020:33:jcs243097. doi: 10.1242/jcs.243097.

Lee G, Leech G, Rust MJ, Das M, McGorty RJ, Ross JL, Robertson-Anderson RM. Myosin-driven actin-microtubule networks exhibit self-organized contractile dynamics. Sci Adv 2021:7:eabe4334. doi: 10.1126/sciadv.abe4334

Maia ARR, Zhu X, Miller P, Gu, G, Maiato H, Kaverina, I. Modulation of Golgi-associated microtubule nucleation throughout the cell cycle. Cytoskeleton 2013:70:32–43. doi: 10.1002/cm.21079.

Mentlik AN, Sanborn KB, Holzbaur EL, Orange JS. Rapid lytic granule convergence to the MTOC in natural killer cells is dependent on dynein but not cytolytic commitment. Mol Biol Cell 2010:21:2241–2256. doi: 10.1091/mbc.e09-11-0930

Miller PM, Folkmann AW, Maia A RR, Efimova N, Efimov A, Kaverina I. Golgi-derived CLASP-dependent microtubules control Golgi organization and polarized trafficking in motile cells. Nat. Cell Biol. 2009:11:1069–1080. doi: 10.1038/ncb1920.

Mimori-Kiyosue Y, Grigoriev I, Lansbergen G, Sasaki H, Matsui C, Severin F, Galjart N, Grosveld F, Vorobjev I, Tsukita S, Akhmanova A. CLASP1 and CLASP2 bind to EB1 and regulate microtubule plus-end dynamics at the cell cortex. J Cell Biol 2005:168:141–153. doi: 10.1083/jcb.200405094.

Ong ST, Chalasani MLS, Fazil MHU, Prasannan P, Kizhakeyil A, Wright GD, Kelleher D, Verma NK. Centrosome-and Golgi-localized protein kinase N-associated protein serves as a docking platform for protein kinase A signaling and microtubule nucleation in migrating T-cells. Front Immunol 2018:397. doi: 10.3389/fimmu.2018.00397

Ori-McKenney K., Jan LY, Jan YN. Golgi outposts shape dendrite morphology by functioning as sites of acentrosomal microtubule nucleation in neurons. Neuron 2012:76:921–930. doi: 10.1016/j.neuron.2012.10.008.

Pariani AP, Almada E, Hidalgo F, Borini-Etichetti C, Vena R, Marín L., Larocca MC. Identification of a novel mechanism for LFA-1 organization during NK cytolytic response. J Cell Physiol 2023:238:227–241. doi: 10.1002/jcp.30921.

Parry RV, Rumbley CA, Vandenberghe LH, June CH, Riley JL.. CD28 and inducible costimulatory protein Src homology 2 binding domains show distinct regulation of phosphatidylinositol 3-kinase, Bcl-xL, and IL-2 expression in primary human CD4 T lymphocytes. JImmunol. 2003:171:166–174. doi: 10.4049/jimmunol.171.1.166

Rak GD, Mace EM, Banerjee PP, Svitkina T, Orange JS.). Natural killer cell lytic granule secretion occurs through a pervasive actin network at the immune synapse. PLoS Biol:2011:9: e1001151. doi: 10.1371/journal.pbio.1001151.

Raemer PC, Kohl K, Watzl C. Statins inhibit NK-cell cytotoxicity by interfering with LFA-1-mediated conjugate formation. Eur J Immunol 2009:39:1456–1465. doi: 10.1002/eji.200838863

Rivero S, Cardenas J, Bornens M, Rios RM. Microtubule nucleation at the cis-side of the Golgi apparatus requires AKAP450 and GM130. EMBO J 2009:28(: 1016–1028. doi: 10.1038/emboj.2009.47.

Roubin R, Acquaviva C, Chevrier V, Sedjaï F, Zyss D, Birnbaum D, Rosnet O.Myomegalin is necessary for the formation of centrosomal and Golgi-derived microtubules. Biol open 2012:2: 238–250. doi: 10.1242/bio.20123392.

Sanders AA, & Kaverina I. Nucleation and dynamics of Golgi-derived microtubules. Front Neurosci. 2015:9:431. doi: 10.3389/fnins.2015.00431.

Sanders AAWM, Chang K, Zhu X, Thoppil RJ, Holmes WR, Kaverina I. Nonrandom γ-TuNA-dependent spatial pattern of microtubule nucleation at the Golgi. Mol Biol Cell. 2017:28:3181–3192. doi: 10.1091/mbc.E17-06-0425.

Schmidt PH, Dransfield DT, Claudio JO, Hawley RG, Trotter KW, Milgram SL, Goldenring JR. AKAP350, a multiply spliced protein kinase A-anchoring protein associated with centrosomes. J Biol Chem. 1999:274:3055–66. doi: 10.1074/jbc.274.5.3055.

Stehbens SJ, Paszek M, Pemble H, Ettinger A, Gierke S, Wittmann T. CLASPs link focal-adhesion-associated microtubule capture to localized exocytosis and adhesion site turnover. Nat Cell Biol. 2014: 16: 558–570. doi: 10.1038/ncb2975.

Takahashi M, Yamagiwa A, Nishimura T, Mukai H, Ono Y. Centrosomal proteins CG-NAP and kendrin provide microtubule nucleation sites by anchoring γ-tubulin ring complex. Mol Biol Cell. 2002:13:3235–3245. doi: 10.1091/mbc.e02-02-0112.

Tonucci FM, Ferretti A, Almada E, Cribb P, Vena R, Hidalgo F, Favre C, Tyska MJ, Kaverina I, Larocca MC. Microtubules regulate brush border formation. J Cell Physiol 2018:233:1468–1480. doi: 10.1002/jcp.26033.

Trambas CM, Griffiths GM. Delivering the kiss of death. Nat Immunol. 2003:4:399–403. doi: 10.1038/ni0503-399.

Vinogradova T, Miller PM, Kaverina I. Microtubule network asymmetry in motile cells: role of Golgi-derived array. Cell Cycle. 2009 Jul 15;8(14):2168–74. doi: 10.4161/cc.8.14.9074.

Wu, J., & Akhmanova, A. (2017). Microtubule-organizing centers. Annual review of cell and developmental biology, 33(1), 51–75.

Yadav, S., & Linstedt, A. D. (2011). Golgi positioning. Cold Spring Harbor perspectives in biology, 3(5), a005322.

Zhu, X, Kaverina I. Golgi as an MTOC: making microtubules for its own good. Histochem Cell Biol. 2013:140:361–7. doi: 10.1007/s00418-013-1119-4.

